# MARK2/Par1b present at retraction fibres corrects spindle off-centering induced by actin disassembly

**DOI:** 10.1101/508861

**Authors:** Madeleine Hart, Ihsan Zulkipli, Roshan Lal Shrestha, David Dang, Duccio Conti, Isabela Kujawiak, Viji Draviam

## Abstract

Tissue maintenance requires adequate cell proliferation and a directed plane of cell division. Retraction fibres can determine the plane of cell division by directing spindle movements; however, retraction fibre components that direct spindle movements remain unclear. We report MARK2/Par1b kinase as a novel component of actin-rich retraction fibres, important for directed spindle movements. A kinase-dead mutant of MARK2 reveals MARK2’s ability to monitor actin status. MARK2’s localisation at retraction fibres, but not the rest of the cortical membrane or centrosome, is dependent on its kinase activity, highlighting a specialised spatial regulation of MARK2. By subtly perturbing the actin cytoskeleton, we demonstrate MARK2’s role in correcting spindle off-centering, induced by lesions in actin assembly. In addition to this mitotic role, we show MARK2’s post-mitotic role in ensuring normal G1-S progression and cell proliferation. We propose that MARK2 provides a molecular framework to integrate cortical signals and cytoskeletal changes in both mitosis and interphase.

**Short Summary:** Coordination of cell proliferation and division is important for tissue maintenance. We report a regulated localisation for MARK2 in mitosis and interphase. We demonstrate its mitotic role in correcting spindle positioning defects and its interphase role in G1-S transition.

## Introduction

Tissue development and regeneration requires a close coordination of cell proliferation, cell division and cell polarity regulatory pathways. With key regulatory function in these pathways, the PAR1/MARK is a family of kinases evolutionarily conserved from yeasts to humans (reviewed in (Wu and Griffin, 2017; Goldstein and Macara, 2007). Dysregulation of the PAR1/MARK signalling has been implicated in a variety of human pathologies including neurodegenerative disorders, carcinomas, and metabolic diseases, making them attractive therapeutic targets (reviewed in Wu and Griffin, 2017).

Among the four human Par1 kinases (MARK 1-4), MARK2/hPar1b is uniquely important for establishing the plane of division and it achieves this through two modes: a cell polarity dependent mode and also a cell polarity independent but cell shape determined mode (Slim et al., 2013; Zulkipli et al., 2018; Lázaro-Diéguez et al., 2013). In both cases, MARK2 controls the position of the mitotic spindle, as seen in a variety of non-polarised and polarised systems including, human cervical epithelial cell cultures, hepatocyte lumina and columnar epithelia (Slim et al., 2013; Zulkipli et al., 2018; Lázaro-Diéguez et al., 2013; Tabler et al., 2010). While cell polarity pathways control spindle positioning by asymmetric enrichment of cortical Dynein, how interphase cell shape determines spindle positioning in non-polarised cultures is not fully clear (reviewed in (Chin et al., 2014; Michel and Dahmann, 2016)).

Retraction fibres and adherens junctions retain the memory of interphase cell shape and tension (Mitchison, 1992; Whur et al., 1977; Harris, 1973; Bergstralh et al., 2017; den Elzen et al., 2009). Altering the distribution of retraction fibres using substrate micropatterning or laser surgery demonstrated the importance of retraction fibres in positioning the spindle (Théry et al., 2007, 2005; Fink et al., 2011). Retraction fibres contain a variety of specialised membrane, cytoskeletal and matrix-adhesion proteins (Mitchison, 1992; Matsumura et al., 2016; Lock et al., 2018; Kwon et al., 2015). However, the nature of signalling events at retraction fibres which are important for spindle positioning remain unclear, hampering our molecular understanding of how retraction fibres determine the plane of cell division.

In polarised epithelia, MARK2/Par1b localises along the basolateral membrane and mediates the development of membrane domains (Suzuki et al., 2004). It’s membrane localisation is thought to be negatively regulated by aPKC mediated phosphorylation at T595 (Suzuki et al., 2004; Hurov et al., 2007) or T508 (Göransson et al., 2006). In addition, MARK2 bears a conserved KA1 domain, which in MARK1 and MARK3 is capable of directing the kinase to specific membrane patches (Moravcevic et al., 2010). Whether MARK2 has multiple modes of interaction at the plasma membrane, and whether its membrane localisation is responsive to cytoskeletal changes needs to be determined to explain how the kinase monitors and regulates spindle positions.

Here, we show how MARK2’s membrane localisation is dynamically regulated in mitotic and interphase cells using several live-cell imaging techniques: Fluorescence recovery after photobleaching (FRAP) and Total Internal Reflection Fluorescence (TIRF) imaging in millisecond time-scales along with long-term time-lapse microscopy. We report a novel localisation for MARK2 kinase at actin-rich mitotic retraction fibres, which is dependent on its kinase activity. Using a kinase dead mutant of MARK2, we uncover a novel role for MARK2 in monitoring cortical actin stress fibres in interphase. We show that MARK2 is required to recenter spindles that are off-centered following actin disassembly, showing the close functional relationship between MARK2 and the actin network. Finally, we report an interphase role for MARK2 in ensuring G1 progression, highlighting MARK2’s role in cell proliferation. We propose that during both interphase and mitosis, MARK2 localises at specialised membrane subdomains and coordinates actin and microtubule cytoskeletal changes, thus enabling normal cell proliferation and division.

## Results

### MARK2 enriches at membrane domains sensitive to actin status

During interphase MARK2 localises along the basolateral membrane in polarised epithelial cells (Suzuki et al., 2004), but it’s underlying regulation is not fully understood. To study whether MARK2’s activity can influence its membrane localisation, we generated HeLa FRT/TO cell lines that conditionally express either YFP fused MARK2-Wild-type (WT) or one of two MARK2 point mutants: MARK2 D157A (kinase dead mutant; KD) or MARK2 T595E (a point mutant to mimic aPKC phosphorylation proposed to block its membrane localisation; (Hurov et al., 2004)) (Fig. 1A). Controlled expression of all three forms of YFP-fused MARK2 (Wild-type, KD and T595E mutant) could be achieved following Doxycycline treatment (Fig. 1B). Live-cell imaging of interphase HeLa FRT/TO cells expressing YFP-MARK2 showed that both the kinase dead mutant and the T595E point mutant enrich at the plasma membrane similar to MARK2-wild-type protein (Fig. 1C). Comparing deconvolved single-plane images of 3D-stacks showed YFP-MARK2 localisation as discontinuous punctate patches at the cell-substrate interface (Z_CS_), and a relatively continuous cortical membrane localisation in deeper Z-slices (Z_3_; 1.5 microns above the substrate) confirming membrane localisation in in all 3 forms of MARK2. Thus, the phosphorylation of MARK2 at T595 by aPKC is insufficient to displace MARK2 from the plasma membrane.

**Fig. 1.**
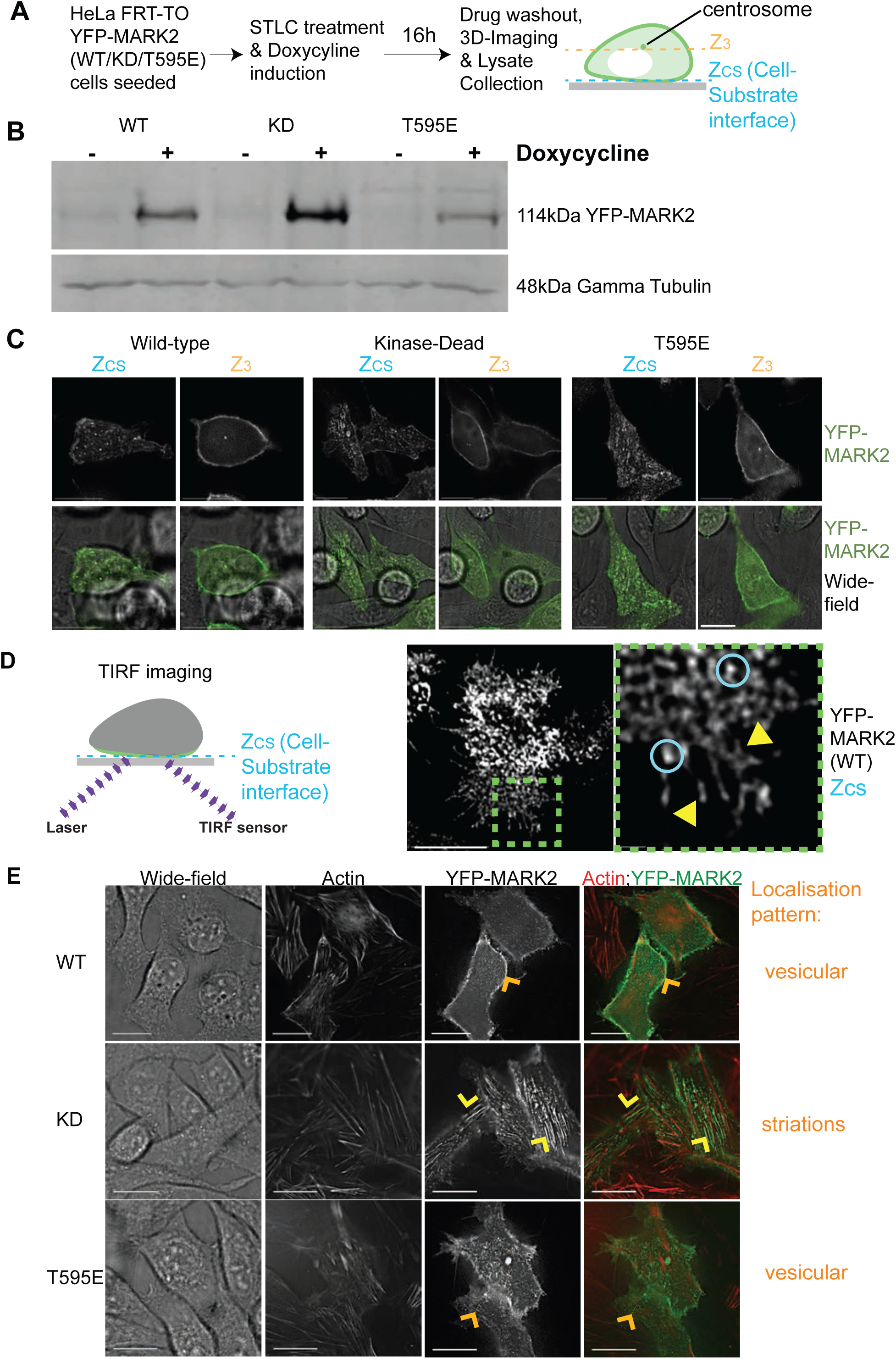
MARK2 membrane localisation is independent of its kinase activity and Thr595 phosphorylation status. (**A**) Experimental Regime: Expression of YFP-MARK2 (WT; wild-type) or YFP-MARK2 (KD; kinase dead) or YFP-MARK2 (T595E) mutants was induced in HeLa FRT/TO cell lines using Doxycycline. To observe MARK2 localisation in interphase and mitosis, YFP-MARK2 expressing HeLa FRT/TO cells were enriched for mitosis using STLC, an Eg5 inhibitor, that blocks monopolar to bipolar spindle conversion. Cells were imaged soon after STLC washout. **(B)** Immunoblot showing the conditional expression of YFP-MARK2 (WT), YFP-MARK2 (KD) or YFP-MARK2 (T595E) mutant in HeLa FRT/TO cell lines upon Doxycyline treatment. **(C)** Representative Z-sections from 3D image stacks showing cell-substrate and cell cortex localisation of YFP-MARK2 (WT), YFP-MARK2 (KD) or YFP-MARK2 (T595E) mutant as indicated. ‘Wide-field’ refers to white light images acquired to indicate the periphery of rounded up mitotic cell. **(D)** Cartoon illustrates TIRF microscopy of YFP-MARK2 at cell substrate interface. Images in the right (uncropped and cropped as indicated) show YFP-MARK2 localisation as submicron sized vesicles (blue rings) or membrane patches (yellow arrows) that move through time (rest of the time-lapse images see Supplementary Figure-1 and Supplementary Movie). Scale: 10µm and insets 1µm. Cells were treated with Doxycyline and Control SiRNA 48h before imaging. **(E)** Deconvolved Z-slice of 3D image-stacks show YFP-MARK2 membrane patches at the cell-substrate interface in WT, KD or T595E mutant expressing cells, as indicated, stained with SiRActin dye. Yellow arrow indicates dense membrane patches of YFP-MARK2-KD signal as ‘striations’ observed along Actin stress fibres. Orange arrows indicate sparse vesicular YFP-MARK2 arranged parallel to actin stress fibres in WT and T595E mutant expressing cells. Scale: 5µm

To study the regulation of YFP-MARK2 localisation at the plasma membrane, we performed Total Internal Reflection Fluorescence (TIRF) imaging using a Super-resolution microscope OMX-SR^TM^. YFP-MARK2 (WT) is present at the plasma membrane as dynamic submicron sized domains (Fig.-1D; see Supplementary Fig.-1 and Supplementary movie for time-lapse images). The MARK2 kinase dead (KD) mutant localised as prominent long striations aligned along actin stress fibres stained using sirActin dye (Fig. 1E), suggesting MARK2’s ability to monitor cortical actin status. Consistent with this notion, YFP-MARK2 signals in Wild-type and T595E mutant cells were observed as vesicles densely arranged parallel to the orientation of the actin stress fibres. In summary, these live-cell studies demonstrate a highly dynamic and regulated membrane localisation for MARK2 at the cell-substrate interface in interphase: first, MARK2 enriches at the plasma membrane as dynamically moving submicron-sized patches, and second, its membrane localisation pattern adjacent to actin fibres (but not membrane association) is dependent on its kinase activity.

### MARK2 is a retraction fibre component regulated by its activity

During mitosis MARK2 is needed for the correct spindle positioning (Zulkipli et al., 2018) and the distribution of retraction fibres can influence spindle positioning(Théry et al., 2005). Hence, we tested whether MARK2 is enriched at retraction fibres found at the cell-substrate interface. We enriched for mitotic cells (Fig. 2A) and analysed YFP-MARK2 localisation at both the mid-cortex region (where spindle poles are visible) and at the cell-substrate interface (where retraction fibres are visible) (Fig. 2A). Both the wild-type (WT) and the two point mutants of YFP-MARK2 (KD and T595E) were normally enriched at the spindle poles and cortical membrane corresponding to the mid-cortex region, as expected (Fig. 2B). However, MARK2 membrane localisation at the cell-substrate interface was dissimilar between the WT and KD mutant. Deconvolved Z-slices corresponding to the cell-substrate interface (Zcs) showed patches of YFP-MARK2-WT signals along retraction fibres that extend out of the rounded-up mitotic cell (Fig. 2C), revealing a novel mitotic localisation for MARK2 at the cell-substrate interface. Importantly, YFP-MARK2 signal along the retraction fibres is reduced in length in cells expressing the kinase-dead mutant, but not the T595E mutant or the Wild-type kinase (Fig. 2C & 2D), indicating a role for MARK2’s kinase activity in regulating its levels at the retraction fibres. The proportion of cells without or with (long & short) YFP-signal at retraction fibres did not differ significantly between cells expressing the wild-type or mutant forms of MARK2 (Fig. 2E). Taken together, the data show that the membrane localisation of MARK2 along retraction fibres, but not the rest of the cortical membrane, is dependent on its kinase activity.

**Fig. 2.**
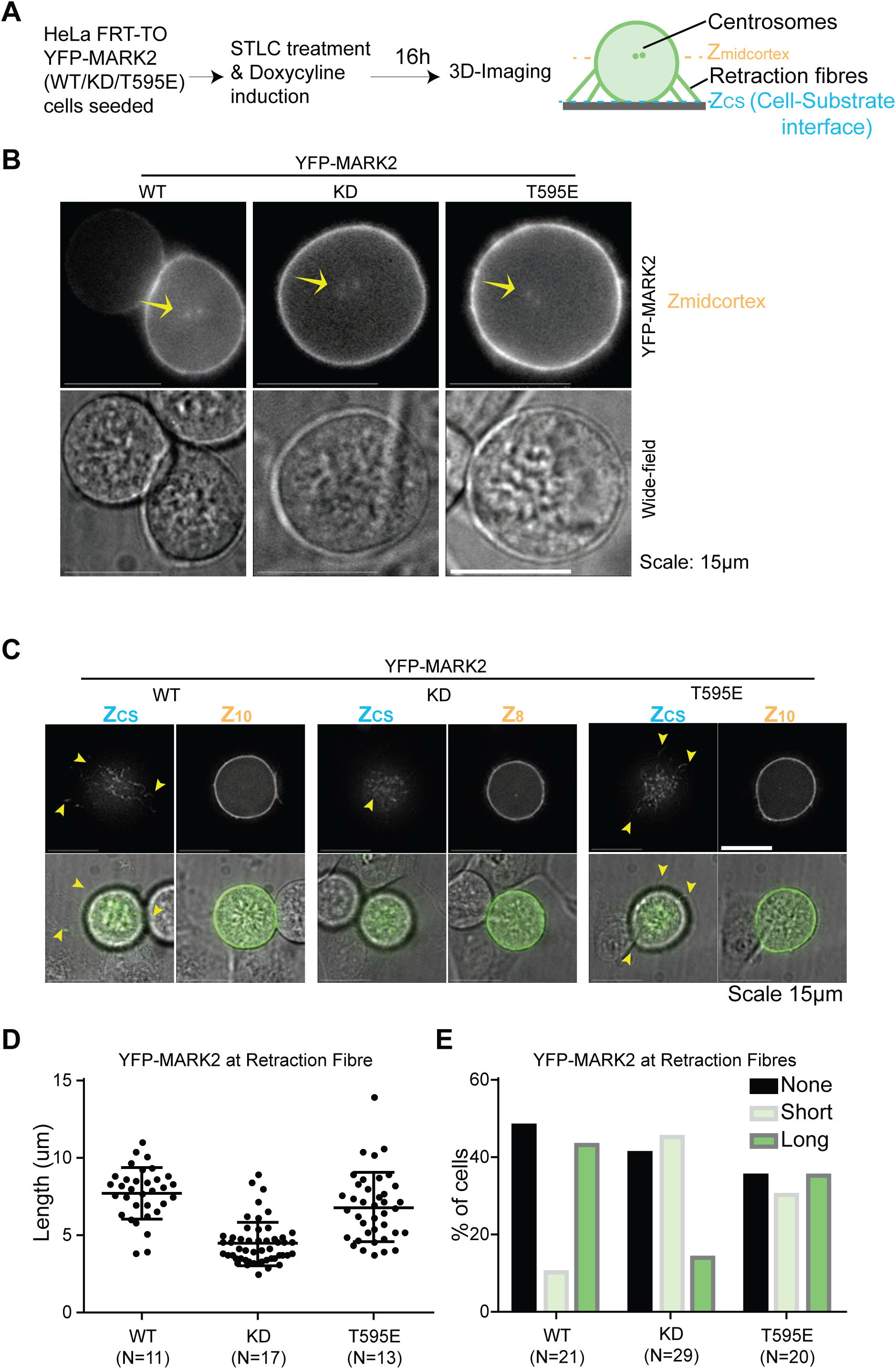
MARK2 localisation at retraction fibres, but not the rest of the cortex or centrosomes, depends on its activity. (**A**) Experimental Regime: Expression of YFP-MARK2 (WT; wild-type) or YFP-MARK2 (KD; kinase dead) or YFP-MARK2 (T595E) mutants was induced in HeLa FRT/TO cell lines using Doxycycline. To observe MARK2 localisation in mitotic cells, YFP-MARK2 expressing HeLa FRT/TO cells were enriched in prometaphase of mitosis using STLC, an Eg5 inhibitor, that blocks monopolar to bipolar spindle conversion. Cells were imaged soon after STLC washout. **(B)** Representative live-cell still images of HeLa FRT/TO cells expressing YFP-MARK2 (WT), YFP-MARK2 (KD) or YFP-MARK2 (T595E) mutant as indicated. Widefield refers to white light images acquired to indicate the periphery of rounded up mitotic cell. Yellow arrows refer to separating spindle poles soon after STLC washout. **(C)** Representative Z-sections from 3D image stacks showing cell-substrate and cell cortex localisation of YFP-MARK2 (WT), YFP-MARK2 (KD) or YFP-MARK2 (T595E) mutant as indicated. Widefield refers to white light images acquired to indicate the periphery of rounded up mitotic cell. Yellow arrows refer to retraction fibres at cell-substrate or cell-cell interface. **(D)** Graph showing the distribution of YFP signal lengths at retraction fibres in HeLa FRT/TO cells expressing one of the three forms of MARK2 as indicated. N refers to number of mitotic cells. **(E)** Graph of percentage of cells showing the presence (long and short) or absence of YFP-MARK2 at retraction fibres assessed using YFP signals in HeLa FRT/TO cells expressing one of the three forms of MARK2 as indicated. N refers to number of mitotic cells.

We investigated whether the centrosome localisation of MARK2 is dependent on its activity. To address this, HeLa cells coexpressing MARK2 KD-YFP and RFP-PACT (RFP fused to Pericentrin PACT domain) were treated with MG132 for 60 minutes to enrich for metaphase cells. MARK2 KD-YFP colocalised with the RFP-PACT signal, indicating that the mutant can be recruited to the centrosome (Supplementary Fig. 2A). To exclude the possibility of indirect enrichment of MARK2-KD at centrosomes due to vesicular traffic towards spindle poles, cells were incubated with 1.7 μM Nocodazole for 30 minutes to depolymerise all microtubules. The loss of microtubules and the bipolar spindle structure were inferred from the random location of centrosomes in metaphase. MARK2 KD-YFP localises normally at the centrosomes of nocodazole treated cells (Supplementary Fig. 2B). Therefore, neither the activity of MARK2 nor microtubules is required to localise MARK2 at the centrosome, highlighting MARK2’s instrinsic ability to bind to centrosomes. Thus, MARK2 enrichment at retraction fibres alone, but not the rest of the cortical membrane or centrosomes is regulated by its kinase activity.

### MARK2 is recruited independent of cortical dynein or microtubules

We investigated whether the cortical membrane localisation of MARK2 is sensitive to cortical Dynein or astral microtubule status. In *C elegans*, microtubule binding of PAR2 provides a protected environment to load PAR2 at the cortex which in turn enriches PAR1 kinase at the cortex (Motegi et al., 2011). Cortical Dynein can capture astral microtubules laterally or end-on (Hendricks et al., 2010; Samora et al., 2011; Laan et al., 2012). Whether cortical dynein or astral microtubules influence MARK2 localisation is not known.

To investigate whether cortical Dynein regulates MARK2 enrichment at cortex, we depleted the cortical Dynein adaptor, LGN using LGN siRNA oligos that we had previously standardised (Corrigan et al., 2013). In LGN siRNA treated cells, YFP-MARK2 was normally localised at the cortical membrane as in control siRNA treated cells, suggesting that cortical Dynein status is not important for MARK2 enrichment at cortex.

To assess whether MARK2 enrichment is sensitive to microtubules, we used Fluorescence Recovery After Photobleaching (FRAP), to bleach a small area of YFP-MARK2 at the cortical membrane and measure YFP-MARK2 recovery, in the presence and absence of microtubules by treating cells either with DMSO (solvent control) or Nocodazole, respectively. FRAP studies showed that MARK2’s membrane localisation is highly dynamic with a fluorescence recovery rate of 7.07 seconds (Fig. 3E) There was no significant change in the rate of YFP-MARK2 recovery in Nocodazole treated or untreated cells (Fig. 3E), showing that astral microtubules do not regulate MARK2 dynamics at the cortical membrane.

**Fig. 3.**
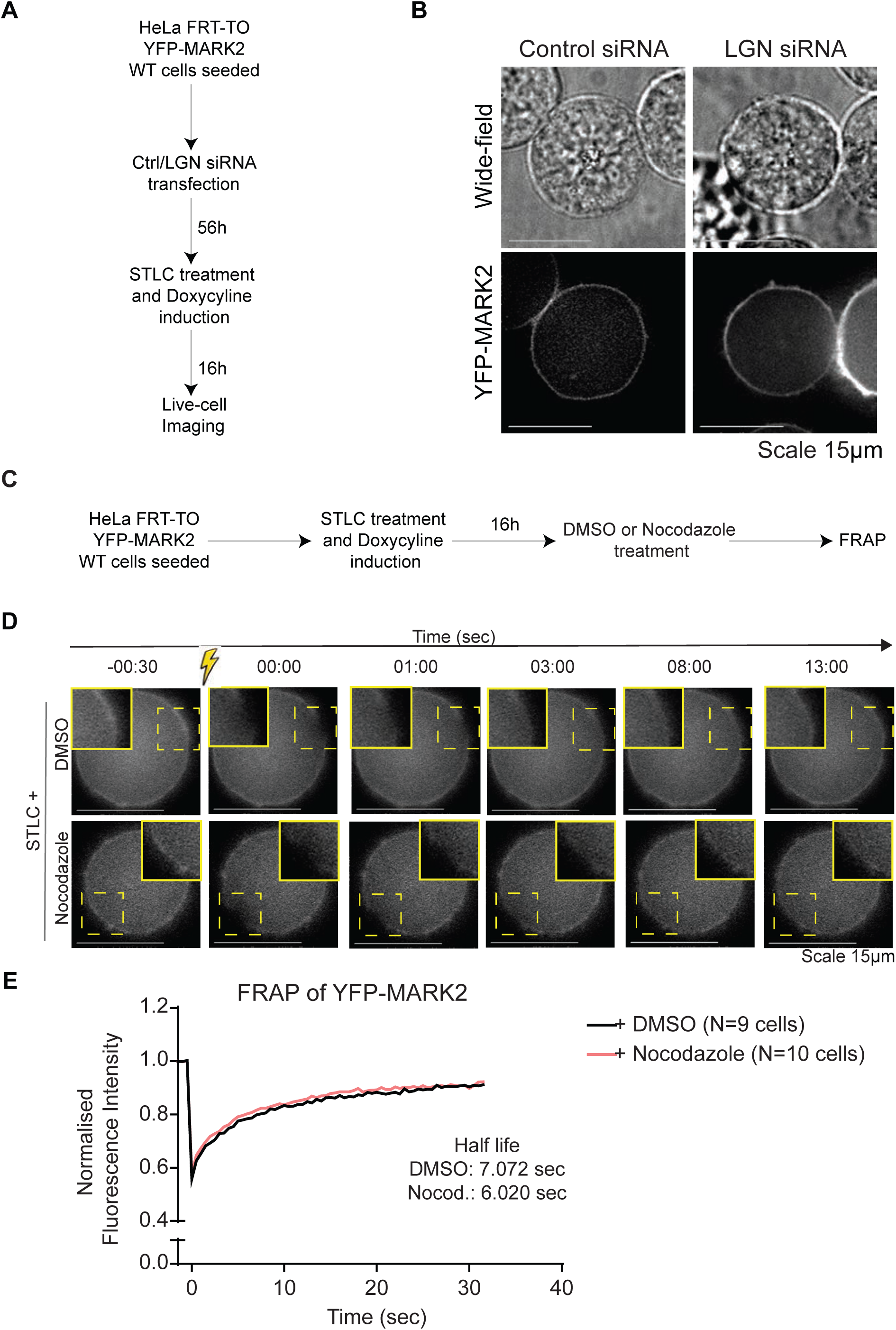
MARK2 localisation at the mitotic cortex is highly dynamic and independent of cortical Dynein or microtubules. (A) Experimental regime: HeLa FRT-TO YFP-MARK2 cells were transfected with siRNA and induced to express YFP-MARK2 using Doxycyline. Cells were treated with STLC to enrich for mitotic cells in prometaphase. **(B)** Representative live-cell images of HeLa FRT-TO cells expressing YFP-MARK2 WT that were transfected with Control or LGN siRNA 72 hours prior to filming. Z-slices corresponding to mid-cortex are shown. Wide-field transmission channel and YFP channel are shown. Scale bar: 15µm. **(C)** Experimental Regime - HeLa FRT-TO YFP-MARK2 Wild type cells were seeded. YFP-MARK2 expression was induced with Doxycyclin and cells arrested at pro-metaphase using STLC. 16 hours later cells were treated with STLC and either DMSO or Nocodazole (as indicated) was added prior to Fluorescence Recovery After Photobleaching (FRAP). **(D)** Images of cells (treated as in C) before and after FRAP. Yellow square indicates bleach area. **(E)** Quantification of FRAP intensities in cells treated as in A and shown in B. Background values were subtracted and values were normalised to opposing cortical values, as well as pre-bleach values (collated from two sets). Each cell was bleached at 3 cortical areas and recovery of fluorescence taken at each site. FRAP rates were obtained Non-linear fit plateau followed by one phase decay.

The FRAP studies and LGN depletion studies demonstrate that the cortical enrichment of MARK2 is insensitive to the status of astral microtubules and cortical Dynein - two established key regulators of spindle movements. These findings position MARK2 as an independent upstream regulator of the spindle positioning process.

#### MARK2 but not Dynein is enriched at actin-rich retraction fibres

To compare the localisation of cortical Dynein and MARK2 kinase, we performed Deconvolution live-cell microscopy of SiR-Actin stained HeLa FRT/TO cells expressing YFP-MARK2 (WT) and HeLa cells expressing Dynein Heavy Chain fused to GFP (DHC-GFP). As expected, mitotic cells expressing YFP-MARK2 showed a clear localisation of YFP-MARK2 and SiR-Actin signals along the retraction fibres in Z-slices close to the cell-substrate interface (Fig. 4A). At higher Z-slices, away from the cell-substrate interface, cortical YFP-MARK2 signal was frequently observed along the outer surface of cortical SiR-Actin signal (Fig. 4A; left), consistent with MARK2’s membrane localisation. Unlike YFP-MARK2, no DHC-GFP signal was observed at retraction fibres in HeLa cells expressing DHC-GFP. At higher Z-sections, where the spindle signal of DHC-GFP was visible, cortical DHC-GFP signal was frequently observed along the inner surface of cortical actin signal (Fig. 4A; right). Based on the 3D-image stacks, we conclude that MARK2 but not Dynein is found at the retraction fibres, and that cortical Dynein and MARK2 occupy inner and outer layers of the cell cortex, respectively. (Fig. 4B).

**Fig. 4.**
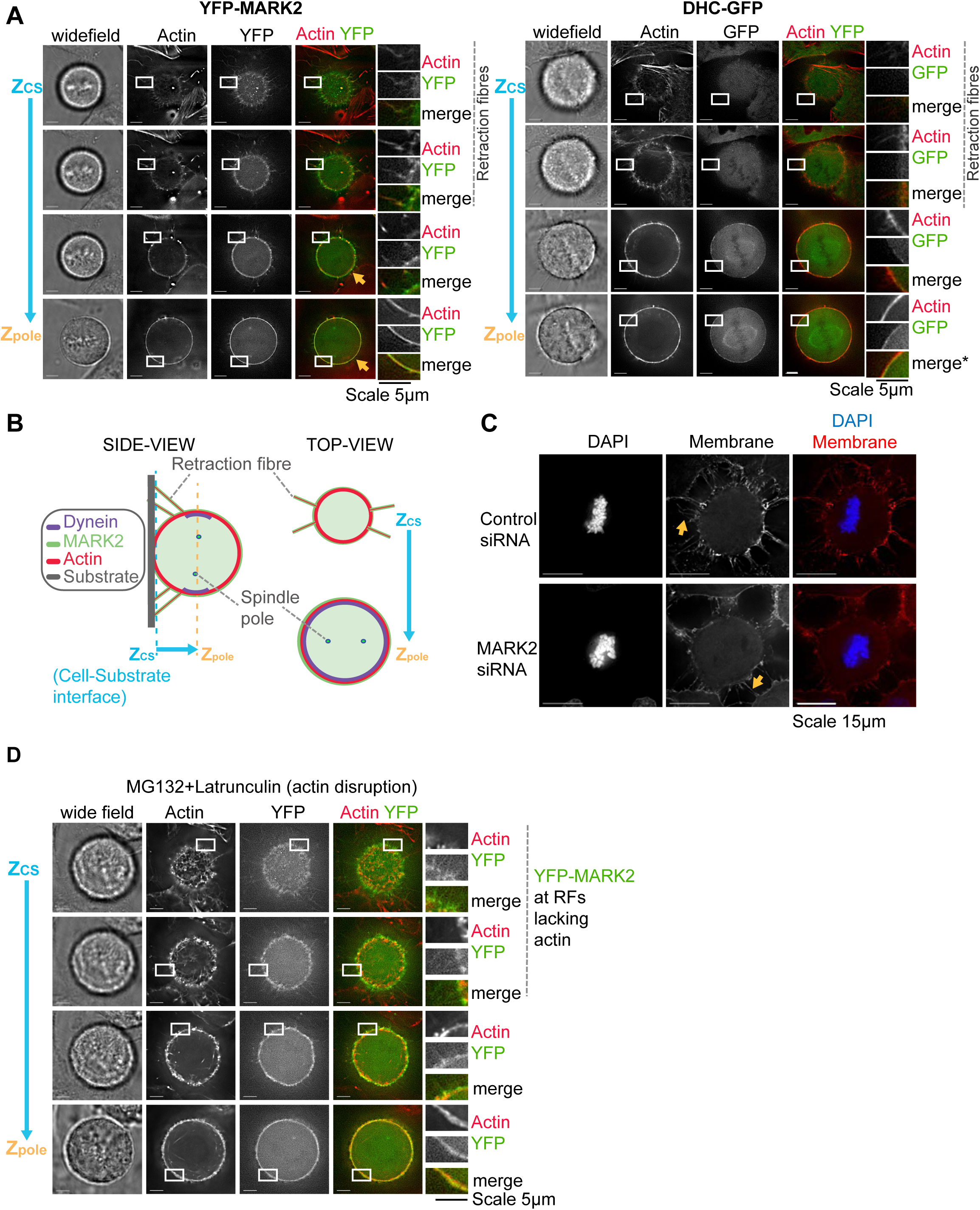
MARK2 and Dynein occupy distinct areas of the mitotic cell cortex. (**A**) Live-cell deconvolved single-plane images of a Doxycyline treated HeLa FRT/TO cell expressing YFP-MARK WT (left) and a HeLa cell expressing mouse DHC-GFP from an endogenous promoter (right) stained with SiR-actin dye for 30 minutes prior to imaging. Two Z-slices of 3D-image stacks close to Cell-substrate interface (Zcs) show the presence of YFP-MARK2 signal (left) and the absence of DHC-GFP signal (right) at retraction fibres marked using SiR-Actin dye (in red). Z-slices of 3D-image stacks near the spindle pole position (Z_pole_) show YFP-MARK2 signal (in green) enveloping cortical actin signal (in red; left images, marked using yellow arrows) and cortical actin signal enveloping DHC-GFP signal (in green; right images, merge crop marked *). **(B)** Cartoon showing the relative positioning of Z-slices (i) at Cell-substrate interface (Z_cs_) where retraction fibres are visible and (ii) above (Z_pole_) where the two spindle poles are visible. Both side-view and top-view of 3D-image stacks presented. Top view images illustrate the presence of Cortical Dynein signal at Z_pole_ but not Z_CS_. **(C)** MARK2 is not essential for the formation of retraction fibres: Representative images of cells treated with siRNA, as indicated were fixed and stained with CellBrite^™^ (membrane marker) and DAPI (DNA marker). Single-plane Images shown are Z-slices corresponding to cell-substrate interface extracted from deconvolved 3D-image stacks. Orange arrows mark SiR-membrane stained retraction fibres that stretch out of the rounded-up cell cortex. Scale bar: 15µm **(D)** MARK2 localisation at retraction fibres is independent of actin: Live-cell images of Doxycyline treated HeLa FRT/TO cells expressing YFP-MARK (WT) exposed to Latrunculin-A and SiR-actin dye for 30 minutes prior to imaging. Z-slices of 3D-image stacks corresponding to spindle pole position show YFP-MARK2 signal enveloping cortical actin signal and Z-slices of 3D-image stacks corresponding to cell-substrate interface show YFP-MARK2 signal (in green) at retraction fibres lacking SiR-actin signal (in red). Scale bars: 5µm.

### Retraction fibres form independent of MARK2

MARK2 is recruited to retraction fibres in a kinase activity dependent manner; whether MARK2 is needed for the formation or maintenance of retraction fibres is not known. To address this, we assessed retraction fibres using a membrane marker, CellBrite. To deplete MARK2 previously standardised siRNA oligonucleotides against MARK2 were used (Zulkipli et al., 2018). Both Control siRNA treated cells and MARK2 siRNA treated cells displayed membrane signal corresponding to retraction fibres at the cell-cell interface (Fig. 4C). We conclude MARK2 is not essential for the formation or maintenance of mitotic retraction fibres.

As retraction fibres form normally in the absence of MARK2, we investigated whether the disruption of actin will affect MARK2 localisation at Retraction Fibres. To disrupt actin in metaphase, we treated cells with low doses of Latrunculin and MG132 (a proteasome inhibitor that blocks anaphase onset) and studied the localisation of MARK2 in retraction fibres that lacked SiR-Actin signals. In Retraction fibres that lacked SiR-Actin we did not observe any stark reduction in YFP-MARK2 signal, indicating that YFP-MARK2 recruitment at retraction fibres is independent of actin status at retraction fibres.

### MARK2 corrects spindle off-centering caused by actin disassembly

MARK2 regulates microtubule length and it ensures normal equatorial centering of spindles (Zulkipli et al., 2018). In addition to microtubules, an intact actin network and proper crosslinking of actin filaments to microtubules are also required for normal positioning of spindles (Gachet et al., 2001; Kunda et al., 2008; Solinet et al., 2013; Kwon et al., 2015).

Because MARK2 localises independently of actin at retraction fibres, we asked whether actin disassembly induced spindle off-centering is monitored and corrected by MARK2. To address this, we studied equatorial spindle off-centering and re-centering in cells treated with low doses of Latrunculin in the presence and absence of MARK2. Spindle off-centering and recentering events were monitored in HeLa coexpressing mCherry-Tubulin and Histone 2B-GFP using time-lapse images acquired once every four minutes (Corrigan et al., 2013). As expected, time-lapse movies of cells treated with Control siRNA, but not MARK2 siRNA, showed rapid recentering of equatorially off-centered spindles within 4-8 minutes (Fig. 5A). We next quantified the percentage of off-centered to centered (OC to C) correction episodes within a 8 minute time window. Nearly 73% of Control siRNA treated cells showed at least 50% success in correcting off-centered spindles (OC to C episodes within 8 min.). However, only 29% of MARK2 siRNA treated cells showed at least 50% success in correcting off-centered spindles (OC to C episodes within 8 min), although spindle recentering at anaphase was nearly 80% (Supplementary Fig. 3B), confirming MARK2’s pre-anaphase role in equatorial centering of spindles (Zulkipli et al., 2018). This is consistent with differential control of anaphase and pre-anaphase spindle positioning events (reviewed in (Bergstralh and St Johnston, 2014).

**Fig. 5.**
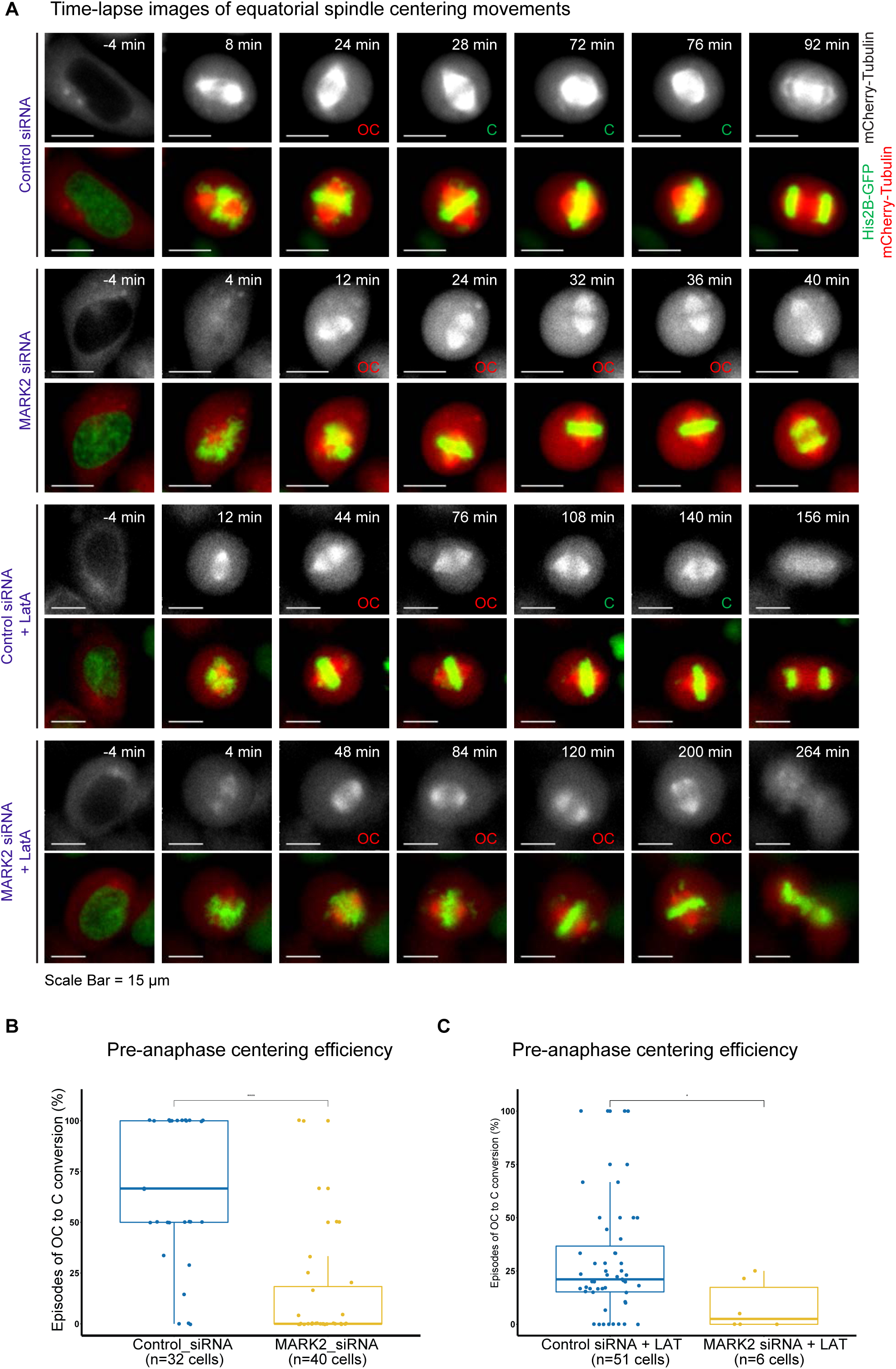
MARK2 is needed to correct spindle off-centering induced by actin disassembly. (**A**) Representative time-lapse image stills of HeLa (His-GFP;mCherry-Tubulin) cells transfected with siRNA as indicated. 1 hour prior to live-cell imaging, the actin perturbant Latrunculin A (Lat-A) was added and time lapse imaging performed for 5 hours. OC refers to off-centered spindles (assessed on the basis of equatorial spindle position within the cell boundary (inferred from mCherry Tubulin signals on the spindle and the cytoplasm). (**B** and **C)** Box plot shows percentage of episodes of equatorially off-centered spindles that centered within an 8-minute time window. For pair-wise comparison a non-parametric Wilcoxon signed-rank test was performed. Asterisks indicate statistical significance for p-value <= 0.05 N values indicate number of cells from 3 independent repeats.

To assess whether MARK2 plays a role in correcting spindle off-centering events induced by actin disassembly, cells were exposed to a low dose of Latrunculin 30 minutes prior to imaging (Supplementary Fig. 3A). Latrunculin treatment induced an increased incidence of spindle off-centering in Control siRNA treated cells (Fig. 5A), showing the need for an intact actin network for proper equatorial centering of spindles. Nevertheless, 25% of Control siRNA treated cells exposed to Latrunculin showed at least 50% success in correcting off-centered spindles (OC to C episodes within 8 min), suggesting an underlying mechanism to monitor and correct spindle off-centering induced by actin disassembly. We next analysed MARK2 siRNA treated cells exposed to Latrunculin: first, the number of cells that initiated mitosis was reduced (n=6 cells, compared to >30 cells in Control siRNA treated cells exposed to Latrunculin). Strikingly, none of the MARK2 siRNA treated cells exposed to Latrunculin showed at least 50% success in OC to C episodes within 8 minutes (Fig. 5C). However, the proportion of off-centered to centered spindles in anaphase is similar in both Control and MARK2 depleted cells exposed to Latrunculin (Supplementary Fig. 3C). We conclude that MARK2 is important for correcting spindle off-centering induced by actin perturbation during pre-anaphase stages of mitosis, a phase when the spindle undergoes rotational movements.

To quantify the efficiency of the correction process, we next compared the rates of OC to C episodes in the presence and absence of MARK2 in Latrunculin treated cells. A cumulative frequency graph showed that 70% of OC to C episodes were completed within either 8 minutes or 16 minutes following Control siRNA or MARK2 siRNA treatment, respectively (Supplementary Fig. 3D), showing a slight delay in OC to C rates following MARK depletion. In stark contrast, OC to C episodes were very delayed in MARK2 siRNA treated cells exposed to Latrunculin: 70% of OC to C episodes were at least 50 minutes long, compared to 16-20 minutes in Control siRNA treated cells. Comparing OC to C rates in Latrunculin treated cells in the presence and absence of MARK2 confirms MARK2’s role in correcting spindle off-centering induced by actin disassembly (Supplementary Fig. 3B). We conclude that MARK2 is essential for correcting spindle off-centering induced by changes in the actin network.

### MARK2 ensures normal progression through G1-phase

In our time-lapse movies of MARK2 siRNA treated cells, we observed a noticeable reduction in the number of mitotic cells which motivated us to explore MARK2’s role in cell cycle progression beyond mitosis. In fact, MARK3, another human homologue of PAR-1a, is required for cell cycle progression and cell proliferation (Müller et al., 2001; Peng et al., 1998). Mouse knockout studies show that MARK2 and MARK3 can compensate for each others’ loss in developing tissues, suggesting that MARK2 may also be important for cell cycle progression and proliferation. To address this directly, we treated HeLa cells with control or MARK2 siRNA and quantified mitotic index 72 hours after siRNA treatment (Fig. 6A). Mitotic index (percentage of mitotic cells) was reduced following MARK2 siRNA treatment compared to control siRNA treatment (Fig. 6B) suggesting a role for MARK2 in ensuring normal cell cycle progression. To confirm MARK2’s cell cycle role, we synchronised cells using Aphidicolin (S-phase block for 24h) following siRNA treatment and then washed Aphidicolin to release cells synchronously into G2 phase (Supplementary Fig 4A). We used two different siRNA oligos against MARK2, standardised in the group (Zulkipli et al., 2018). Time-lapse imaging of cells for a period of 8-13 hours after release from a 24 hour Aphidicolin showed that both siRNAs against MARK2 reduced the number of cells entering mitosis (initiating NEBD) compared to control siRNA treated cells (Supplementary Fig 4B). Quantitative analysis of the number of cells entering mitosis per 1024×1024 pixel image field in the 5 hour time-lapse movie showed that while control cells have on average 25 mitotic cells per movie, MARK2 depleted cells had only 7 and 10 mitotic cells for MARK2-1 and MARK2-2 siRNA treated cells, respectively (Supplementary Fig 4C). We conclude that MARK2 depleted cells show a noticeable decrease in the number of cells entering mitosis, demonstrating MARK2’s role in controlling cell cycle progression.

**Fig. 6.**
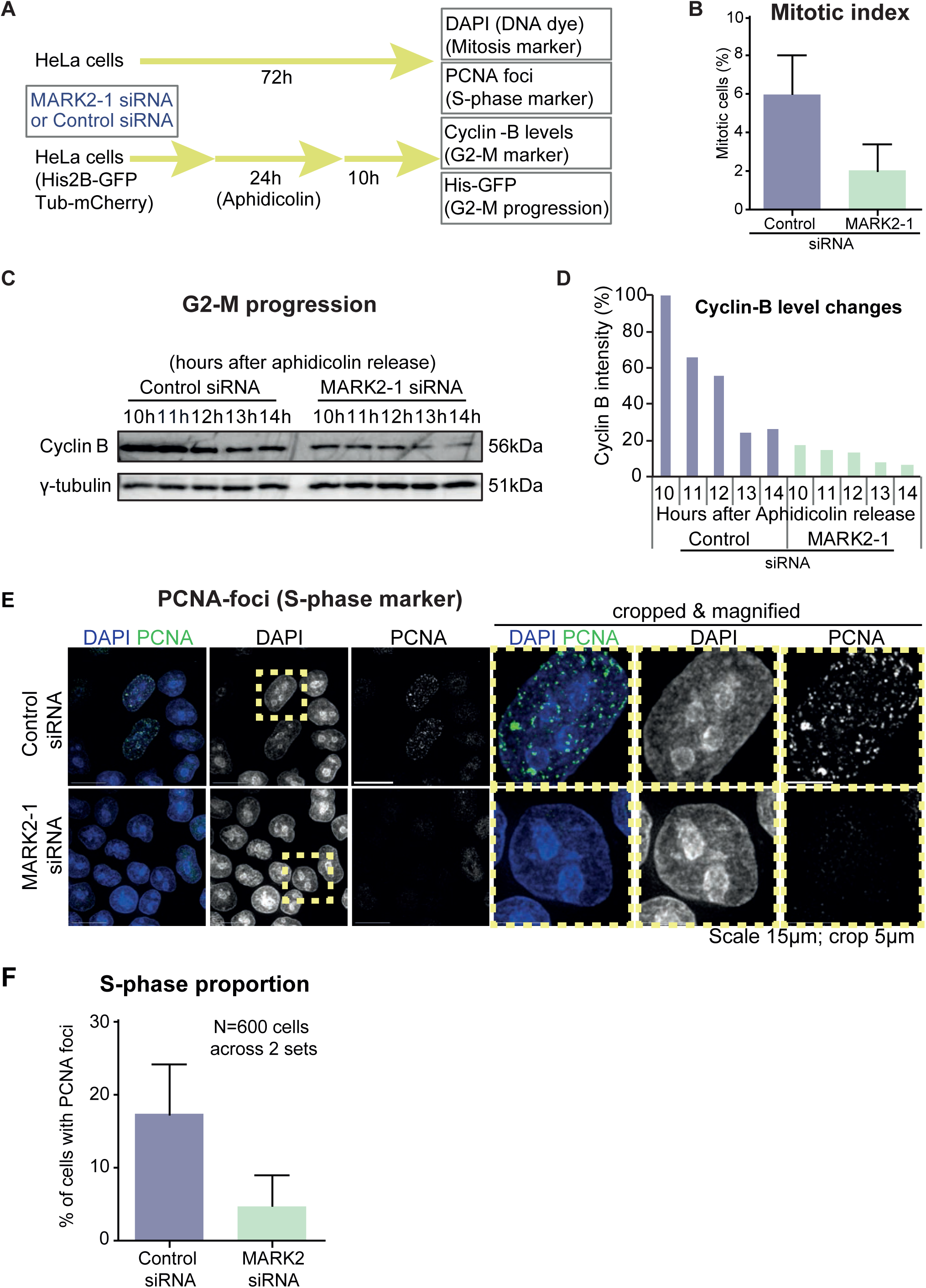
MARK2 ensures normal progression through G1 phase of the cell cycle. (**A**) Experimental Regime: HeLa cell line or Fluorescent HeLa cell line expressing Tubulin-mCherry and Histone 2B-GFP (His2B-GFP and Tub-mCherry) were treated with Control or MARK2 siRNA oligos. 72 hours later, HeLa cells were fixed and immunostained for assessing PCNA foci (S-phase marker) or stained with DAPI for condensed DNA status (mitosis marker). Fluorescent HeLa cells (His2B-GFP and Tub-mCherry) were treated with Aphidicolin for 24 hours for a synchronous release from S-phase and 10 hours later at G2-M boundary assessed every hour for Cyclin B levels. **(B)** Percentage of mitotic cells (with condensed chromosomes) following MARK2 or Control siRNA treated as indicated. Bar graph indicates average values obtained from two independent experiments. Error bars represent SD values calculated and plotted using Prism. **(C)** Immunoblot of cell lysates treated with the indicated siRNA oligos and immunostained with cyclin-B antibody to monitor cell cycle (G2-M) progression after release from a 24 hours aphidicolin block. Gamma-tubulin is used as a loading control. Time indicated is hours after wash-off of aphidicolin. Fluorescent immunoblotting was used to quantify differences in Cyclin-B levels. **(D)** Fluorescent band intensity values corresponding to Cyclin B relative to gamma-tubulin (loading control) were measured as percentage values from 2 independent repeats of experiments as shown in C. Normalised average percentage values are presented as a bar graph. **(E)** Representative images show the presence or absence of PCNA-foci (a marker of DNA replication) in the nuclei of HeLa cells treated with siRNA as indicated. Images shown are overlaid Z-slices to assess PCNA-foci throughout the nuclei. Cropped images are magnified images corresponding to dashed boxes marked in uncropped images. Scale bars in uncropped and cropped images refer to 15 and 5 microM, respectively. **(F)** Bar graphs show the percentage of DAPI-positive nuclei with PCNA foci in cells treated with Control or MARK2 siRNA as indicated. N_cells_ correspond to 600 cells from two independent experiments. Error bars represent SD values calculated and plotted using Prism.

To study whether the reduced mitotic index observed in MARK2-depleted cells is due to a G2 or G1 phase delay, we measured the amount of Cyclin B1, a mitotic cyclin that is high at G2-M transition and destroyed at metaphase-anaphase transition (Clute and Pines, 1999). Immunoblot of control cell lysates showed high levels of Cyclin B1 at 10 hours following aphidicolin release, which as expected, steadily declined through time as cells exited mitosis (Fig. 5C and 5D). In contrast, MARK2 siRNA treated cells do not show a comparable high level of Cyclin B at a similar time point, although tubulin levels were comparable (Fig. 5D), indicating a pre-G2 phase delay in MARK2 depleted cells. To test whether the cell cycle delay in MARK2 siRNA treated cells arose from a delay in G1 or S phase, we investigated the percentage of cells that display PCNA foci, a marker of ongoing DNA replication (Essers et al., 2005). A four fold reduction in the number of PCNA-foci positive cells were observed in cells treated with MARK2 siRNA compared to control siRNA (Fig. 6E and 6F), showing a stark reduction in the number of S-phase cells. In sum, the reduced mitotic index in asynchronous cell cultures, failure to accumulate cyclinB in synchronised cultures, and reduction in PCNA-foci all together demonstrates a role for MARK2 in G1 progression. We conclude that MARK2 ensures timely progression through G1-phase of the cell cycle.

## Discussion

Here we report MARK2/Par1b, a known regulator of microtubule stability, as a novel component of retraction fibres with a role in correcting spindle off-centering induced by actin disassembly. Its recruitment to retraction fibres alone, but not the rest of the mitotic cortical membrane, is dependent on its kinase activity. Importantly, it dynamic localisation at the mitotic cell cortex is independent of cortical Dynein, astral microtubule and actin network, highlighting its upstream position among the regulators of spindle movements. In addition to this mitotic role, MARK2 is also needed for cell proliferation and progression through the G1 phase of the cell cycle. During interphase, MARK2/Par1b is enriched at the interphase plasma membrane as sub-micron sized punctate patches that are highly mobile and coincident with actin stress fibres in a kinase activity dependent manner. We propose that MARK2 recruitment to specialised membrane subdomains can be regulated to monitor and mediate localised cytoskeletal changes both in mitosis and interphase.

MARK2 regulates microtubule dynamics during both interphase and mitosis (Cohen et al., 2004; Nishimura et al., 2012; Mandelkow et al., 2004; Hayashi et al., 2011; Schaar et al., 2004; Zulkipli et al., 2018). We had shown that spindle centering defect observed in MARK2-depleted cells can be rescued by stabilising microtubules (Zulkipli et al., 2018). However, it was unclear how a cortex and spindle pole localised MARK2 can precisely regulate astral microtubule length to achieve equatorially spindle centering. Here, we report that MARK2 localises at retraction fibres in a kinase activity dependent manner. This opens a new paradigm for MARK2 activating/inactivating enzymes (phospho-regulation of MARK2) to locally regulate microtubule dynamics in actin-rich areas (retraction fibres; Figure 7A). Such localised regulation is crucial as microtubule and actin network changes have to be spatially and temporally coordinated during mitosis (reviewed in (Lancaster and Baum, 2014)).

**Fig. 7.**
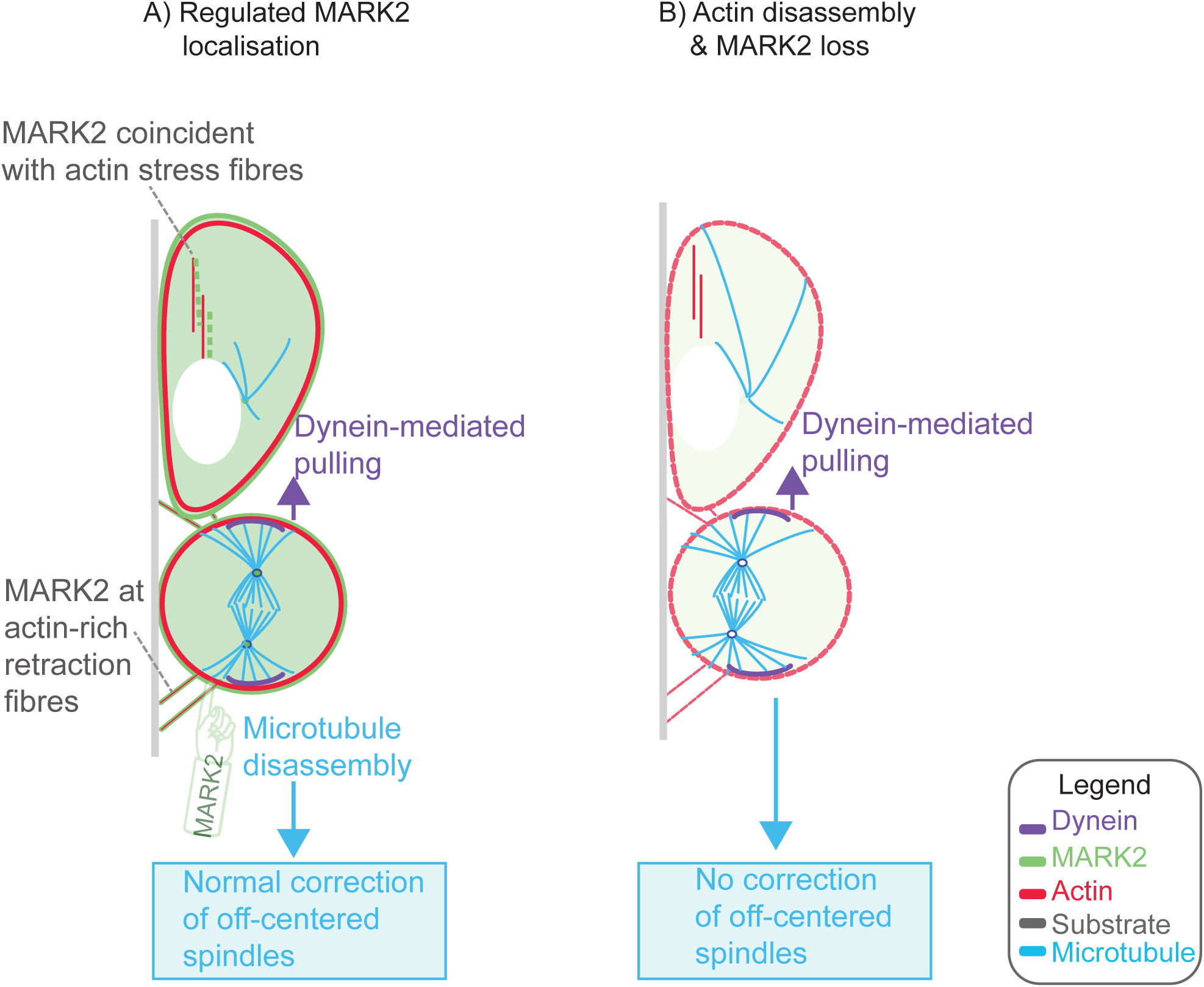
Illustration of MARK2 mediated localised regulation of spindle position. Cortical MARK2 and Dynein occupy outer and inner cortical areas, in alignment with their distinct roles in spindle centering and pulling, respectively. MARK2 can recognise actin status in interphase (A). Actin perturbation induced spindle off-centering can be corrected by MARK2 (B). To enable spindle centering, MARK2 at the cortical membrane can selectively act on astral microtubule ends that reach retraction fibres.

Our findings are consistent with a model where MARK2 dynamically links actin and microtubule cytoskeletal status independent of cortical dynein (see below). MARK2’s localisation along actin stress fibres is regulated by its kinase activity, indicating the kinase can monitor actin status. Moreover, MARK2 mediated phosphorylation of Tau can attenuate Tau binding to F-actin (Cabrales Fontela et al., 2017). We propose that MARK2 at actin-rich retraction fibres is well positioned to correct off-centered spindles by regulating cortical microtubule dynamics (Fig. 7A). In the absence of MARK2, actin disassembly induces spindle off-centering that can not be corrected normally (Fig. 7B). MARK2-mediated spindle recentering is independent of cortical Dynein function for two reasons: first, MARK2 and Dynein Heavy Chain occupy different regions of the cell cortex (Fig. 7A). Second, in the absence of cortical dynein (LGN depletion) neither MARK2 localisation (in this study) nor spindle centering is lost (Zulkipli et al., 2018).

At least one of the four alleles of MARK3 or MARK2 is needed for mouse embryogenesis (Lennerz et al., 2010), suggesting overlapping roles for the two kinases. MARK2 knockout mice are dwarf, infertile and display a variety of complex cellular and tissue-level problems: insulin hypersensitivity, reduced adiposity, reduced learning and memory, and immune system dysfunction (Bessone et al., 1999; Hurov et al., 2007; Segu et al., 2008; Hurov et al., 2001). MARK3 knockout mice show reduced adiposity with normal insulin handling (Lennerz et al., 2010). MARK3 is required for cell cycle progression (Peng et al., 1998; Müller et al., 2001) and here, we show that MARK2 is also required for G1 progression. Thus, the study expands MARK2’s cell cycle role from mitosis (M phase) ((Zulkipli et al., 2018) and this study) to the G1 phase when proliferative fate of cells are determined.

Targeting kinases to specific membrane domains is thought to contribute to “coincidence detection”, a mechanism that allows kinases to mediate events in a localised fashion (Moravcevic et al., 2010). Moreover, spatial exclusivity is known to play an important role in mitotic signalling (Alexander et al., 2011). In the case of MARK2, a coincidence detection mechanism can operate at retraction fibres as MARK2 localisation is controlled by its own kinase activity. Although a single-site phosphorylation at Thr595 (Hurov et al., 2004) is insufficient to block MARK2 recruitment to membrane patches, it is possible that alternate or multiple sites on MARK2 need to be phosphorylated by aPKC for 14-3-3 interaction and its delocalisation from membrane. For example, an alternate site T508 when mutated to Ala increases MARK2’s membrane localisation (Göransson et al., 2006). In the case of xPar-1, xPKC phosphorylates multiple sites - Thr593 and Ser646 (Kusakabe and Nishida, 2004), and in the case of closely related MARK3, aPKC phosphorylation at 17 sites are needed to fully abrogate 14-3-3 interaction (Göransson et al., 2006). Regulating MARK2’s membrane localisation can be an important signalling gateway for external cues to locally control MARK2-mediated cytoskeletal changes during cell division and cell proliferation.

## Materials and Methods

### Cell culture and synchronization

HeLa cells were cultured in DMEM (Tergaonkar et al., 1997) supplemented with 10% FCS and antibiotics (penicillin and streptomycin) and plated onto glass-bottomed dishes (LabTek) or 13-mm round coverslips for imaging. For inhibition experiments, cells were treated with 10 µM MG132 (1748; Tocris). Cells were synchronized using either a single 1 µg/ml aphidicolin block for 24 h and then released for 7 h before filming or by exposing cells to 20µM STLC for 16h prior to imaging. To disassemble microtubules, cells were treated with 1.7 µM nocodazole (Thermo Fisher Scientific).

### Cell line generation

The HeLa^hMARK2-siRES-YFP^ cell line was generated by transfecting a Tet-inducible expression vector encoding siRNA-resistant YFP-hMARK2 and followed by colony picking (Zulkipli et al., 2018). Vectors bearing point mutants of hMARK2 were generated by PCR-based point mutagenesis and confirmed by DNA sequencing. Cell line generation procedures were followed according to the FRT/TO system protocol (Invitrogen). Doxycyclin induction was usually performed for 16h prior to imaging (unless differently specified). 200ng/ml Dox was used in conventional DMEM supplemented with 10% FCS. HeLa DHC-GFP cells line expressing mouse DHC-GFP through an exogenous promoter is a kind gift from the Hyman laboratory. HeLa His2B-GFP mCherry-Tubulin cell line was generated as described in (Shrestha and Draviam, 2013)

### Plasmid and siRNA transfections

HeLa cells were transfected with siRNAs or plasmid vectors as described previously (Shrestha et al., 2014). siRNA oligonucleotides used to deplete MARK2 were used as standardised previously (Zulkipli et al., 2018): MARK2-1 (5’-CCUCCAGAAACUAUUCCGCGAAGUA-3’) and MARK2-2 (5’-UCUUGGAUGCUGAUAUGAACAUCAA-3’), and LGN (Corrigan et al., 2013). siRNA Oligos were purchased from GE Healthcare. Sequences of plasmid vectors are available upon request.

### Live-cell time-lapse imaging and analysis

High Resolution TIRF with deconvolution images were acquired using OMX-SR^™^. Control siRNA treated cells were treated with Doxycyline for 16-20 h prior to imaging. Images were acquired every second for 20 seconds, with 3 z-stacks each 0.125nm apart, and deconvolved. Timepoints were equalised.

FRAP was performed on Deltavision Core^™^ microscope using Quantifiable Laser module components (488 laser). The bleach site was bleached with 488 laser with a pulse duration of 1 second and laser power of 20%. 3 pre-bleach images were acquired 0.5 seconds apart, and 64 post-bleach images were acquired 0.5 seconds apart. Time 0 refers to the first post-bleach image acquired. Fluorescence intensities at the bleach spot and an opposing cortical site were measured using ImageJ. Values were corrected to the ratio between opposing cortical site at each time-point relative to pre-bleach values (to account for acquisition associated photobleaching), and normalised relative to pre-bleach values. Halftimes were calculated using Non-linear regression analysis (plateau followed by one phase decay) using Prism Graphpad.

Cells were transfected with siRNA oligonucleotides or plasmid vectors, 72 or 24 h, respectively (72h for MARK2, Control and LGN siRNA). Before imaging, cells were transferred to Leibovitz L15 medium (Invitrogen) for imaging at 37°C. To observe chromosome and spindle movements, images were acquired with exposures of 0.1 s from at least three Z planes 3 µm apart every 4 min for 5 h using a 40× 0.75 NA objective on an DeltaVision Core^™^ microscope (GE Healthcare) equipped with a Cascade2 camera under EM mode. For colocalising DHC-GFP, YFP-MARK2 and SiR-Actin signals, live-cells were imaged (at least 15 Z-slices, 0.5 µm apart) using a 100× 1.2 NA objective on the microscope described above.

For MARK2 (centrosome) colocalization studies, the centrosome marker (a fragment of pericentrin-tagged to RFP) was used (kind gift from J. Pines, ICR, London). Time-lapse videos were analyzed manually using SoftWoRx^™^. Spindles in HeLaHis-GFP;mCherry-Tub cells were visually scored as equatorially off centered when unequal distances were observed between the cell cortex and the two opposing edges of metaphase plate (histone-GFP signal) or the cell cortex and the two walls of the spindle at the equator (mCherry-tubulin signal) as described in (Zulkipli et al., 2018).

### Immunofluorescence and immunoblotting

For immunofluorescence the PCNA antibody (1:1000, Cell Signalling Technologies; 2586S) was used. For membrane visualisation the membrane dye (Biotium, CellBrite^™^, 30023) was incubated on cells for 15minuted prior to fixation, as per manufacturer’s instructions. DNA was stained with DAPI. Images of immunostained cells were acquired using a 100× 1.2 NA objective on a DeltaVision Core microscope equipped with CoolSnap HQ Camera (Photometrics).

For immunoblotting, antibodies against γ-tubulin (1:800, T6793; Sigma-Aldrich), GFP (1:1000, MBL, 598) and Cyclin B1 (1:1000, BD Bioscience, 554176) were used. Immunoblots were developed using fluorescent secondary antibodies (LI-COR Biosciences), and fluorescent immunoblots were imaged using an Odyssey imager (LI-COR Biosciences).

### STLC wash-off assay

Cells were treated with 20µM STLC for 16h to enrich for mitotic cells. Prior to imaging, media was changed and STLC washed off with at least 4 quick washes in Leibovitz media.

### Latrunculin treatment

Cells were treated with 1µM Latrunculin A with or without 10uM MG132 in Leibovitz media and imaged 30 minutes later.

### Statistical analysis

RStudio Software (v1.1.456) with R distribution (v3.5.1) for statistical computing and GraphPad Prism 5.0 (GraphPad Software, Inc., San Diego, CA) were used to generate graphs and perform statistical analysis. Visual and analytical assessment of normality and homogeneity of variance was performed with QQ-plot and Shapiro-Wilk normality test. The choice for non-parametric tests is in alignment with the underlying data structure. To determine the statistical significance of differences between the population mean ranks in the experimental conditions, a Wilcoxon signed-rank test was used as indicated in the Fig. legends. Box plots show median, upper (75%), and lower (25%) quantiles as the box, whiskers represent 1.5 times the interquartile range, and the remaining points are shown as outliers. All data points were included in statistical analyses.

Following convention for asterisk symbols indicating statistical significance was used: (ns) for p > 0.05, () for p <= 0.05, (*): p <= 0.01, (**\***) p <= 0.001 and (**\*\***) for p <= 0.0001. For OC to C rates in MARK2 and Control siRNA treatments alone data presented in (Zulkipli et al., 2018) was analysed. Error bars show SEM values obtained across experiments or cells as indicated in legend. P-values representing significance were obtained using Mann-Whitney *U* test, proportion test, or paired sample *t* test.

## Acknowledgments

We thank Jorn Breumlund (GE-Healthcare) for technical support while performing TIRF microscopy using OMX-SR^™^, and the Pines group (ICR) for the plasmid vector encoding pericentrin fragment. This work was supported by a Cancer Research UK Career Development Award (C28598/A9787), a Biotechnology and Biological Sciences Research Council Project grant (BB/R01003X/1), and a Queen Mary University of London Laboratory startup grant to V.M. Draviam, a Queen Mary, University of London PhD studentship to M. Hart, a Biotechnology and Biological Sciences Research Council LIDO-DTP studentship to D. Dang and a Universiti Brunei Darussalam PhD studentship to I. Zulkipli.

The authors declare no competing financial interests.

Author contributions: MH, RS, IK and IZ designed and performed experiments and analyzed and interpreted data. MH, RS, IZ, DC and IK analyzed data, and prepared Fig.s. DD supported statistical data analysis and DHC data imaging. VMD planned the study, discussed the experimental design, analyzed and interpreted data, and wrote the manuscript. VMD and MH edited the manuscript.

## Supplementary Figures

**Supplementary Fig. 1.**
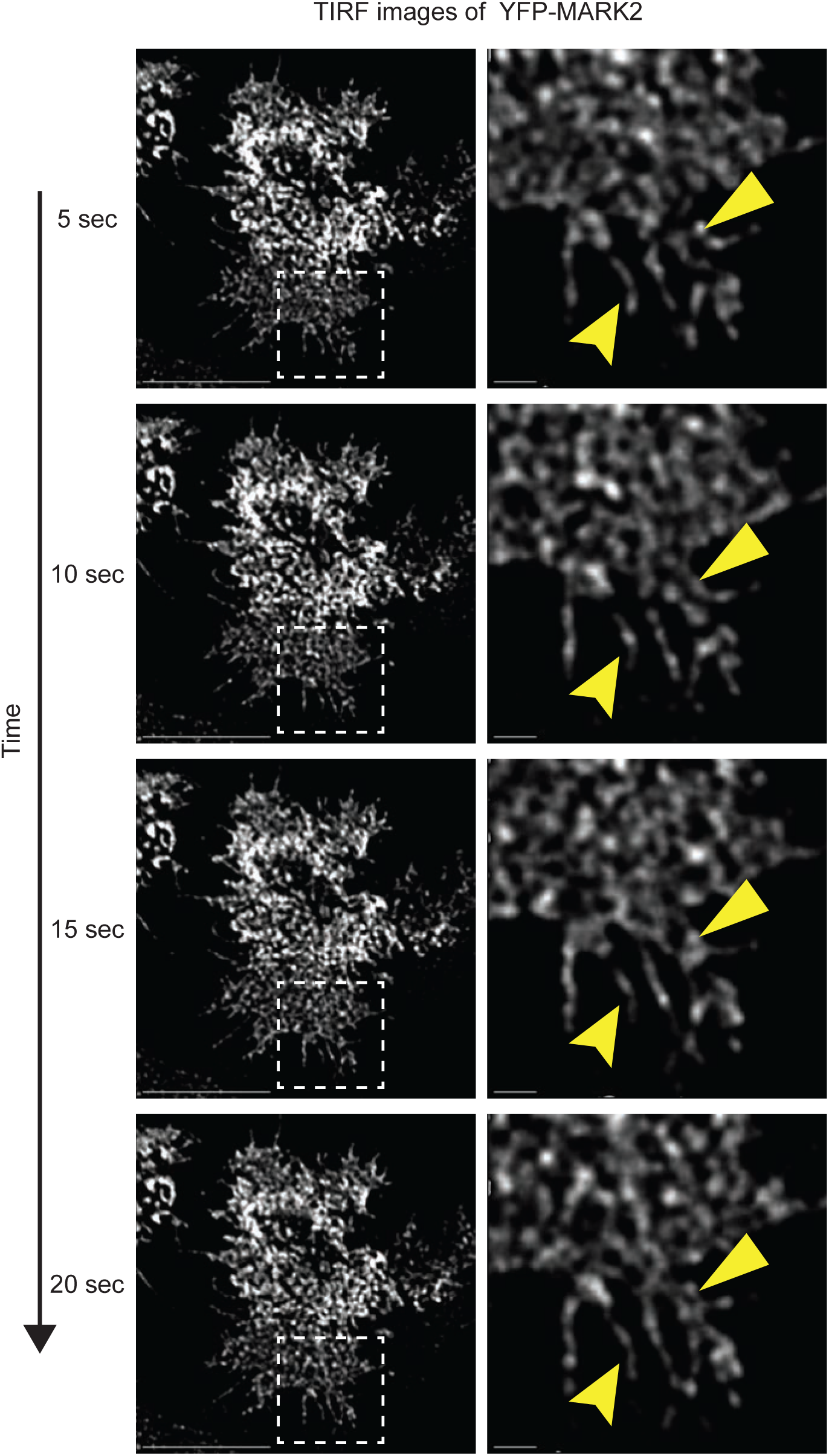
MARK2 is enriched as dynamic membrane patches. Representative still images from a time-lapse TIRF (Total Internal Reflection Fluorescence) microscopy movie of a HeLa FRT-TO cell expressing YFP-MARK2 WT protein. TIRF microscopy images taken once every second show vesicular localisation of YFP-MARK2 and its movement as membrane patches through time (yellow arrows). Time lapse images shown (with time-frames indicated) correspond to the cell in Fig. 1D (Time-frame; 0Sec) and Supplementary Movie. Scale bars: uncropped image, 10µm and cropped image, 1µm.

**Supplementary Fig. 2.**
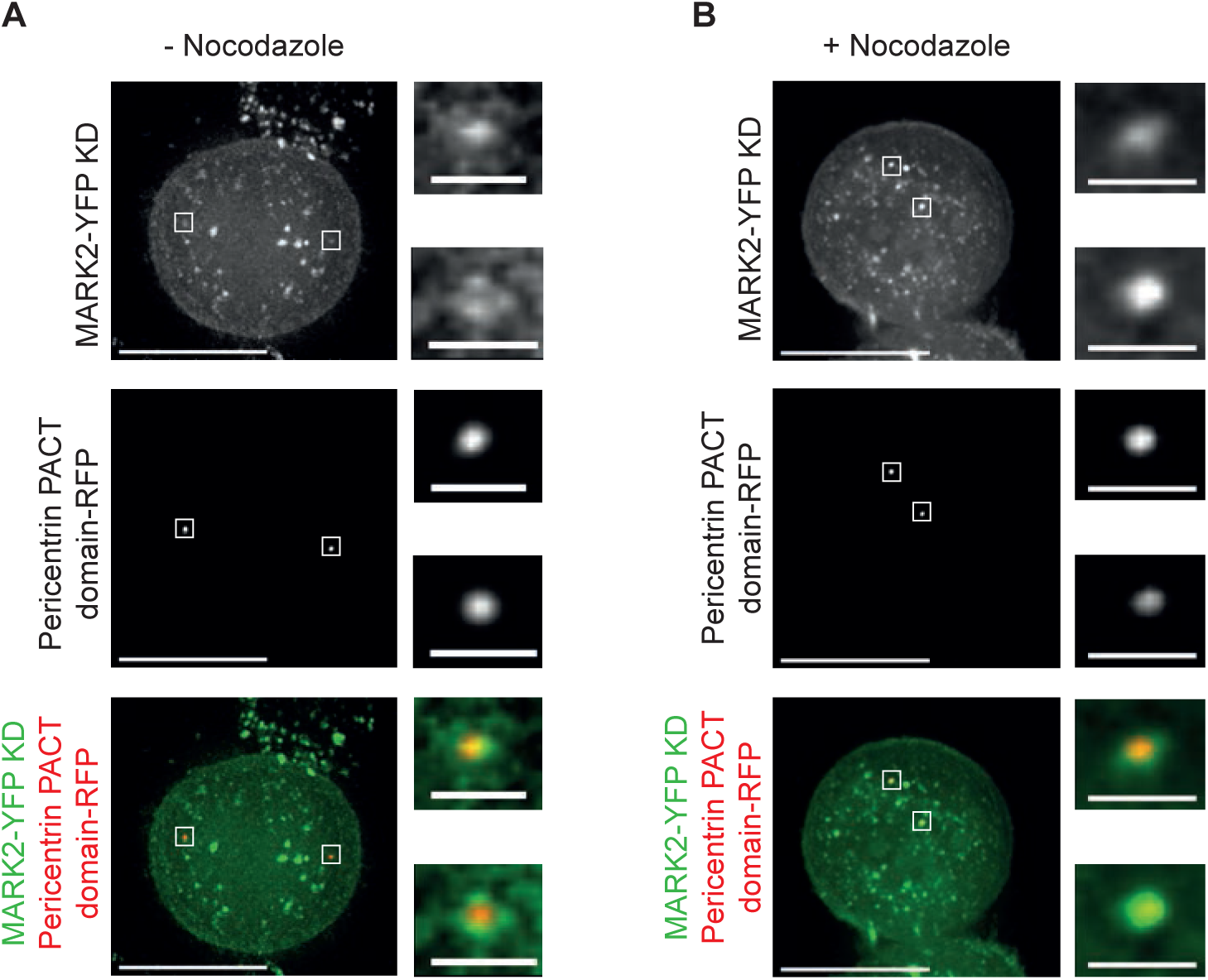
MARK2’s localisation at centrosomes is independent of its activity. Representative live-cell still images of HeLa cells expressing exogenous MARK2-YFP Kinase dead (KD) and the centrosome localising PACT domain of pericentrin tagged with RFP to mark centrosomes, illustrating the localisation of MARK2-YFP KD during (**A**) metaphase and **(B)** metaphase cells exposed to nocodazole treatment. Insets show magnified areas of centrosome as indicated by white boxes on main images. Images of cells in metaphase collected after a 60 minute treatment with MG132. For Nocodazole treatment, cells were exposed to 1.7microM Nocodazole for 30 minutes after MG132 treatment. Scale bars = 10 microns in main image and 1 micron in insets.

**Supplementary Fig. 3.**
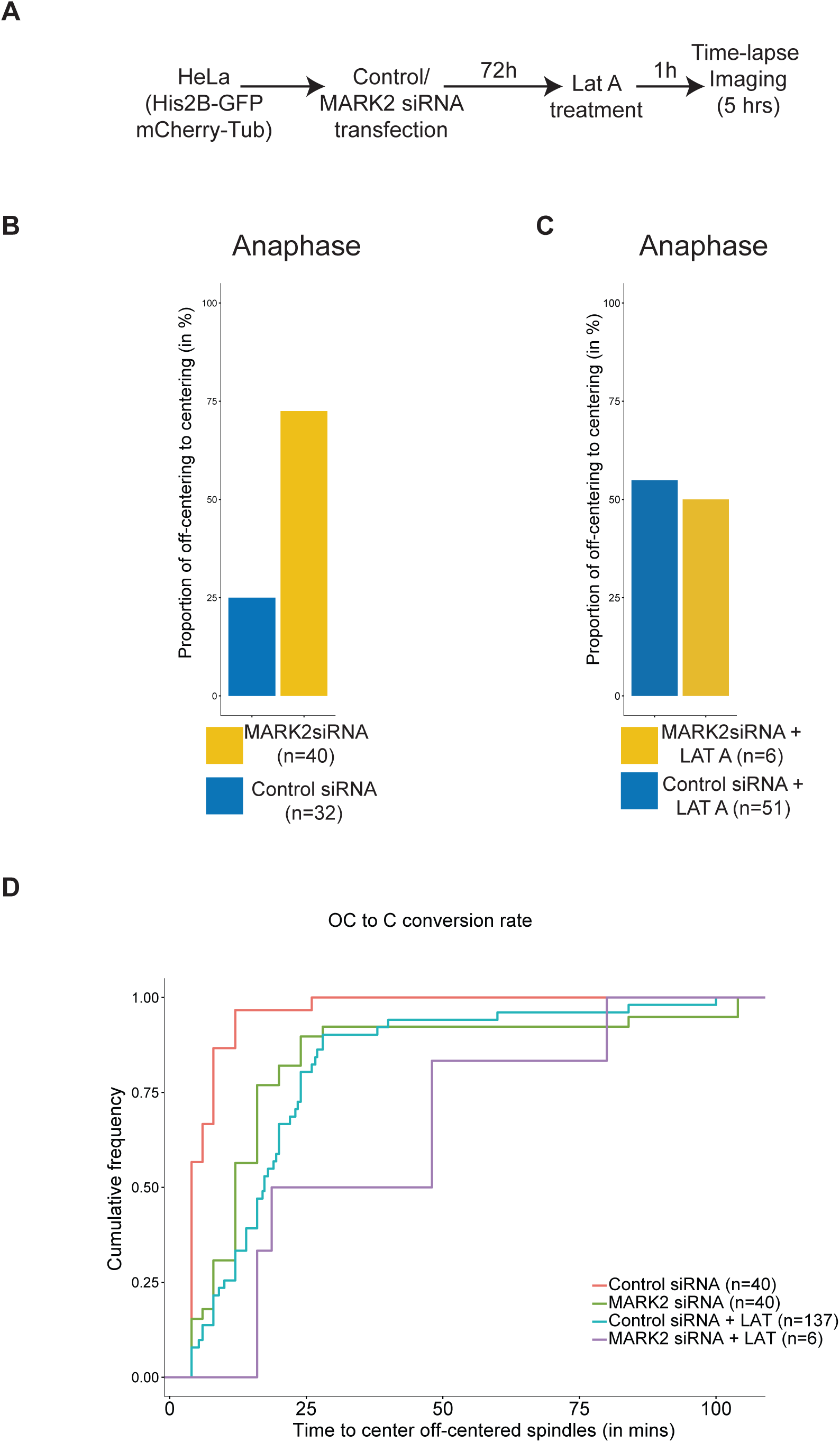
MARK2 enables the recentering of off-centered pre-anaphase spindles. (**A**) Experimental regime: HeLa cells (His-GFP; mCherry-Tub) cells were transfected with Control or MARK2-1 siRNA oligo (as indicated) and 72 hours later imaged once every 4 minutes using a time-lapse Deconvolution microscope. Latrunculin was added 1 hour prior to imaging. Regime corresponds to data show in Fig. 5. **(B and C)** Bar graphs show proportion of cells that centered their spindles at the metaphase-anaphase transition. MARK2 siRNA treated cells center the spindles at metaphase-anaphase transition unlike Control siRNA treated cells that center their spindle pre-anaphase (B). In the presence of Lat-A, cells fail to center spindles even in anaphase (C). n refers to number of cells from at least three independent experimental repeats. **(D)** Empirical Cumulative graph showing the proportion of equatorially Off-Centered (OC) spindles that are successfully Centered (C) within the time (in minutes) indicated. Pre-anaphase and anaphase spindle centering data are all included. Data was obtained from mitotic cells as shown in Fig. 5A.

**Supplementary Fig. 4.**
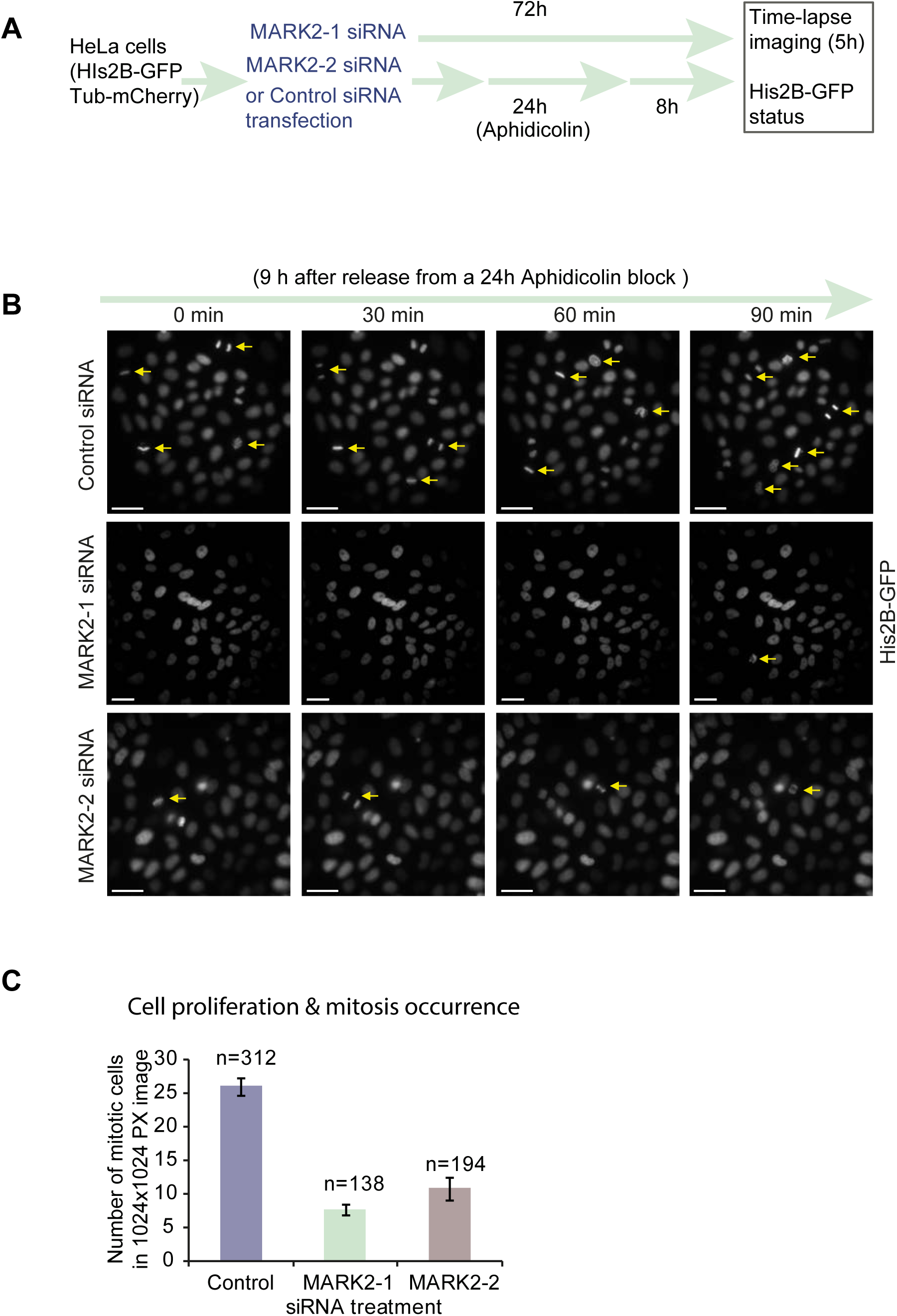
MARK2 depletion reduces cell proliferation and mitosis occurrence. (**A**) Experimental regime: HeLa cells (His-GFP; mCherry-Tub) cells were transfected with Control, MARK2-1 or MARK2-2 siRNA oligos and 72 hours later imaged once every 4 minutes using a time-lapse Deconvolution microscope. 8 hours prior to imaging, cells were released from a 24 hour Aphidicolin treatment to allow synchronous G2-M progression during the time-lapse microscopy session. **(B)** Representative time-lapse still images of HeLa (His-GFP; Tub-mCherry) cells (His-GFP signals shown) transfected with the indicated siRNA oligo and treated as described in (A). Minutes refer to time-lapse image acquisition times (9 hours after release from Aphidicolin; 1 hour after the time-lapse image acquisition was initiated). Yellow arrows indicate cells undergoing mitosis (assessed using His2B-GFP signal of chromosome movements). Scale bar: 30 microns. **(C)** Bar graph shows average number of mitotic cells observed per 1024×1024 pixel image field from a 5 hour long time-lapse movie of cells treated with the indicated siRNA oligo. Total number of cells (n) following various siRNA treatments are indicated. Values for bar graph (mitotic and total number of cells) were obtained from time-lapse movies acquired as in (B). Errors bars represent SEM from 3 independent experiments.

**Supplementary Movie: MARK2 is enriched at mobile membrane subdomains in interphase** Time-lapse TIRF (Total Internal Reflection Fluorescence) microscopy movie of a HeLa FRT-TO cell expressing YFP-MARK2 WT protein. TIRF microscopy images taken once every second show vesicular localisation of YFP-MARK2 and its mobility as membrane patches through time. Time lapse images shown (with time-frames indicated) correspond to the cell in Fig. 1D (Time-frame; 0Sec) and Supplementary Fig. 1.

